# A single amino acid transporter controls the uptake of priming-inducing beta-amino acids and the associated trade-off between induced resistance and plant growth

**DOI:** 10.1101/2022.03.17.484770

**Authors:** Chia-Nan Tao, Will Buswell, Peijun Zhang, Heather Walker, Irene Johnson, Roland Schwarzenbacher, Jurriaan Ton

**Author notes:** corresponding author: Jurriaan Ton. The author(s) responsible for distribution of materials integral to the findings presented in this article in accordance with the policy described in the Instructions for Authors (https://academic.oup.com/plcell/pages/General-Instructions) is : Jurriaan Ton.

## Abstract

Selected beta-amino acids, such as beta-aminobutyric acid (BABA) and R-beta-homoserine (RBH), can prime plants for resistance against broad-spectrum diseases. Here, we describe a genome-wide screen of fully annotated Arabidopsis T-DNA insertion lines for impaired in RBH-induced immunity (*iri*) against the downy mildew pathogen *Hyaloperonospora arabidopsidis*, yielding 104 lines that were partially affected and 4 lines that were completely impaired in RBH-induced resistance. The *iri1-1* mutant phenotype could be confirmed by an independent T-DNA insertion in the same gene, encoding the high-affinity amino acid transporter LHT1. Using uptake experiments with *IRI1/LHT1*-expressing yeast cells and mass spectrometry-based quantification of RBH and BABA in leaves of mutant and over-expression lines of *IRI1/LHT1*, we demonstrate that IRI1/LHT1 acts as the main transporter for cellular uptake and systemic distribution of RBH and BABA. Subsequent characterisation of mutant and over-expression lines of *IRI1/LHT1* for induced resistance and growth responses revealed that the level of *IRI1/LHT1* expression determines the trade-off between induced resistance and plant growth by RBH and BABA.

## INTRODUCTION

The innate immune system enables plants to perceive and react to pathogens and herbivores. The basal component of this regulatory system is under control by pattern recognition receptors that perceive molecular non-self-patterns from the attacker or damaged-self patterns that form during an attack (Choi and Klessig, 2016). Following recognition of these alarm signals, a signalling network is initiated that orchestrates the induction of cellular defence mechanisms, including reactive oxygen species (ROS), callose-rich cell wall depositions and defence-related genes (Chisholm et al., 2006; Bigeard et al., 2015). Apart from this pattern-triggered immunity (PTI), innate immunity can be triggered by susceptibility-inducing pathogen effectors. If the challenged plant expresses a resistance (R) gene that can recognise the activity of such a pathogen effector, the innate immune response is referred to as effector triggered immunity (ETI; Cui et al., 2015). Besides innate immunity, plants can acquire long-lasting resistance, which develops after recovery from biotic stress. This induced resistance (IR) is typically based on priming of the innate immune system, which mediates a faster and/or stronger induction of inducible defences upon secondary attack (Wilkinson et al., 2019; De Kesel et al., 2021). In addition, IR can be triggered by root colonisation of selected plant-beneficial microbes or treatment with specific chemical agents, such as microbe-associated molecular patterns, volatile organic compounds and non-proteinogenic β-amino acids (Mauch-Mani et al., 2017; De Kesel et al., 2021).

β-amino butyric acid-induced resistance (BABA-IR) has emerged as a popular model system to study the molecular mechanisms controlling immune priming in plants. BABA-IR has been reported in more than 40 plant species against different types of pathogens (Cohen, 1994; Cohen et al., 2016). In Arabidopsis (*Arabidopsis thaliana*), BABA primes both salicylic acid (SA) dependent and independent defence mechanisms and protects plants against biotrophic, hemibiotrophic and necrotrophic pathogens (Zimmerli et al., 2000; Ton et al., 2005; Schwarzenbacher et al., 2020). Recent evidence suggests that BABA accumulates during exposure to biotic and abiotic stress (Thevenet et al., 2017), which provides biological relevance and supports previous evidence that an aspartyl tRNA aspartase, IBI1, acts as a plant receptor of BABA (Luna et al., 2014). BABA has also been suggested to act as a microbial rhizosphere signal, based on the finding that induced systemic resistance (ISR) upon root colonisation by *Pseudomonas simiae* WCS417 is blocked in the *ibi1-1* mutant of Arabidopsis (Luna et al., 2014). Despite the apparently high efficiency by which plant roots are capable of taking up BABA from the soil (Zimmerli et al., 2000; Ton et al., 2005), a cellular transporter of this well-known priming agent has never been identified.

Although BABA-IR is effective against a broad spectrum of plant diseases, high doses of BABA results in major growth reduction (Wu et al., 2010; Luna et al., 2014). This undesirable side effect is in part caused by disruptive binding of R-BABA to the aspartic acid-binding pocket of the IBI1 enzyme, causing accumulation of uncharged tRNA^Asp^ and GCN2-dependent inhibition of gene translation (Luna et al., 2014; Buswell et al., 2018). To search for less phytotoxic IR analogues of BABA, we screened a small library of structurally related β-amino acids for IR activity and phytotoxicity in Arabidopsis. This screen resulted in the identification of R-β-homoserine (RBH), which induces resistance in Arabidopsis and the tomato (*Solanum lycopersicum* (L.) cultivar Micro-Tom) against biotrophic and necrotrophic pathogens without growth reductions (Buswell et al., 2018). A recent study comparing four different IR agents for their effectiveness in strawberry (*Fragaria × ananassa*) against *Botrytis cinerea* also identified RBH as the most effective IR agent without negative effects on plant growth (Badmi et al., 2019). Like BABA, RBH primes defence activity of callose-rich papillae, which in Arabidopsis are formed at relatively early stages of infection by the biotrophic Oomycete *Hyaloperonospora arabidopsidis* (*Hpa*). Interestingly, however, despite the structural similarity to BABA, RBH does not require the IBI1 receptor to induce resistance in Arabidopsis (Buswell et al., 2018). Furthermore, unlike BABA, RBH does not prime SA-dependent gene induction but primes for camalexin production upon infection by *Hpa* and jasmonic acid (JA)-dependent defence genes after infection by the necrotrophic fungus *Plectosphaerella cucumerina* (Zimmerli et al., 2000; Ton et al., 2005; Buswell et al., 2018). Hence, RBH-induced resistance (RBH-IR) is controlled by partially different pathways than BABA-IR. So far, the molecular mechanisms responsible for the uptake and perception of RBH have remained unknown.

In this study, we have conducted a genome-wide screen of Arabidopsis T-DNA insertion mutants for impaired in RBH-induced immunity (*iri*) against *Hpa*, yielding 104 and 4 lines that are partially and completely impaired in RBH-IR, respectively. Of the latter, we characterised the *iri1* mutant, which is affected in the high-affinity amino acid transporter LHT1. We provide evidence that the level of *IRI1/LHT1* gene expression determines the balance between IR and plant tolerance by RBH and BABA. Furthermore, mass spectrometry analysis of leaves from RBH- and BABA-treated wildtype, *iri1* and *IRI1*-overexpressing plants revealed that IRI1/LHT1 is critical for the uptake and systemic distribution of both RBH and BABA, while uptake experiments with *IRI1/LHT1*-expressing yeast cells demonstrated that IRI1/LHT1 acts as a high-affinity transporter of BABA and RBH. In support of other studies that have linked IRI1/LHT1 to plant-microbe interactions and plant immunity, we conclude that IRI1/LHT1 acts as a master regulator of the trade-off balance between growth and IR by priming-inducing beta-amino acids.

## RESULTS

### Genome-wide screen for *impaired in RBH-immunity* (*iri*) mutants

To search for new regulatory genes of R-β-homoserine-induced resistance, we screened 23,547 T-DNA insertion lines from SALK and SAIL collection (Alonso and Ecker, 2006) for an impaired in RBH-induced immunity (*iri*) phenotype against *Hpa*, which covers >90% of all annotated protein-coding genes in the Arabidopsis genome. In contrast to conventional EMS-based mutant screens, which rely on selection of mutant phenotypes by individual plants, the collection of fully annotated homozygous T-DNA insertion mutants allowed us to screen small populations of 5 seedlings per line for quantitative *iri* mutant phenotypes, including partial loss of RBH-IR. To reduce false positives, the screen was performed over three successive stages. In the first stage, seedlings were screened in 400-well trays, which were soil-drenched with RBH to a final soil concentration of ~0.5 mM, inoculated with *Hpa* conidiospores and scored for visual sporulation from 5-7 days post inoculation (dpi; Fig. 1A). Each tray yielded ~1-2 lines displaying sporulation in at least 2 seedlings/well by 7 dpi, which were selected and re-screened at stage 2, using the same 400-well tray selection system. Stage 2 yielded a total of 427 putative *iri* mutant lines (Fig. 1A). These putative *iri* mutant lines are taken forward for final validation in stage 3, which was based on categorical scoring of *Hpa* colonisation in trypan-blue-stained leaves from control- and RBH-treated plants (0.5 mM) of each candidate line (Fig. 1A). To validate the statistical robustness of this screening stage, we conducted a pilot experiment that compared *Hpa* colonisation between 40 pots of Col-0 seedlings pre-treated with either water or RBH (0.5 mM). Categorical scoring of trypan blue-stained leaves confirmed statistically uniform distributions of *Hpa* colonisation within each treatment (Supplemental Fig. 1A). Of the 427 putative *iri* lines from stage 2, 104 lines were confirmed to be partially impaired RBH-IR in stage 3, as evidenced by statistically enhanced levels of *Hpa* colonisation in RBH-treated mutant plants compared to RBH-treated wild-type plants, while still showing statistically significant reductions in *Hpa* colonisation by RBH treatment compared to the water controls within each line (Fig. 1A, Supplemental Fig. 1B and Supplemental Data set 1). An additional 4 lines, named *iri1-1* to *iri4-1*, were fully impaired in RBH-IR, as evidenced by statistically identical levels of *Hpa* colonisation between RBH- and water-treated plants within each line (Figs. 1A, Supplemental Fig. 1B and Supplemental Data Set 1).

**Figure 1.**
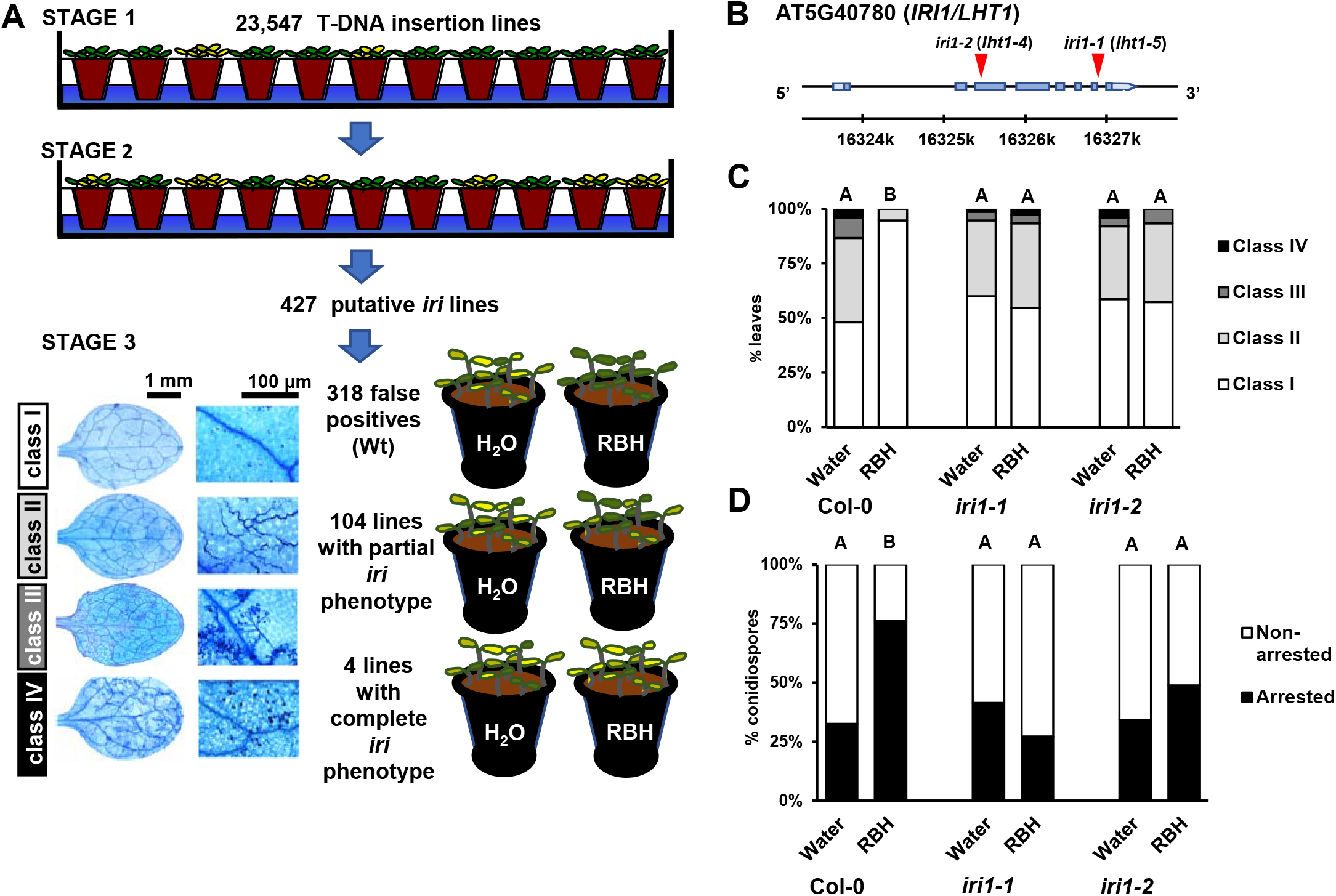
Mutant screen for *<u>i</u>mpaired in <u>R</u>BH-induced immunity* (*iri*) phenotypes and characterisation of the *iri1* mutant of Arabidopsis. **(A)** Scheme of the 3 successive selection stages of *iri* mutant screen of 23,547 T-DNA insertion lines from the SALK/SAIL collection. Small populations of ~5 seedlings per line were screened (stage 1) and re-screened (stage 2) for sporulation by *Hyalopoeronospora arabidopsidis* WACO9 (*Hpa*) upon soil-drenching treatment with 0.5 mM R-β-homoserine (RBH) and subsequent inoculation with *Hpa* conidionspores (top). Putative *iri* lines were validated in controlled RBH-induced resistance (RBH-IR) assays by categorising leaves from water- and RBH-treated (0.5 mM) plants into four *Hpa* colonisation classes at 5-7 days post inoculation (dpi; bottom; Supplemental Fig. 1). Photos of trypan-blue leaves on the bottom right indicate the *Hpa* colonisation classes, ranging from healthy leaves (I), hyphal colonisation with without conidiospores (II), hyphal colonisation with conidiophores (III) to extensive hyphal colonisation with conidiophores and deposition of asexual oospores (IV). **(B)** Gene model of the *IRI1* gene (At5g40780) encoding the Lysine Histidine Transporter1 protein (LHT1). Red triangles indicate two independent T-DNA insertions in the *iri1-1/lht1-5* and *iri1-2/lht1-4* mutants, respectively, to confirm involvement of the *IRI1/LHT1* gene in RBH-IR against *Hpa*. (**C**) Quantification of RBH-IR against *Hpa* in leaves of Col-0, *iri1-1/lht1-5* and *iri1-2/lht1-4*. Shown are frequency distributions of trypan-blue-stained leaves across the four *Hpa* colonisation classes (see **A**). Different letters indicate statistically significant differences between samples at 6 dpi (Fisher’s exact tests + Bonferroni FDR; p < 0.05; n = 70-80 leaves). (**D**) Quantification of arrested *Hpa* colonisation by callose. *Hpa*-induced callose was analysed in aniline blue/calcofluor-stained leaves by epifluorescence microscopy. Shown are percentages of callose-arrested and non-arrested conidiospores at 3 dpi, as detailed by Schwarzenbacher et al. (2020). Different letters indicate statistically significant differences in frequencies between samples (Fisher’s exact tests + Bonferroni FDR; *p* < 0.05; *n* > 100 conidiospores).

### Identification of the *IRI1/LHT1* gene as a critical regulator of RBH-IR against *Hpa*

Since SALK/SAIL lines can carry multiple T-DNA-induced mutations (Alonso and Ecker, 2006), it is possible that the *iri* mutant phenotypes are caused by other gene mutations than those identified and annotated by PCR border recovery analysis. To address this possibility, we quantified RBH-IR in independent T-DNA insertion lines in the annotated genes of each the 4 complete *iri* lines (Figs. 1B-C and Supplemental Figs. 2A-B). Since RBH-IR against *Hpa* in Arabidopsis is associated with augmented effectiveness of callose-rich papillae (Buswell et al., 2018), we quantified the effectiveness callose-mediated cell wall defence at 3 dpi, as detailed previously (Schwarzenbacher et al., 2020). All original *iri* lines consistently lacked RBH-IR and concomitantly failed to augment callose-mediated defence upon RBH treatment (Fig. 1D, Supplemental Fig. 2C), confirming the importance of this post-invasive defence barrier in RBH-IR against *Hpa*. However, independent T-DNA insertions in the annotated genes of *iri2-1, iri3-1* and *iri4-1* did not affect RBH-IR and showed wild-type levels callose-mediated defence against *Hpa* (Supplemental Fig. 2C), indicating that their *iri* phenotypes are caused by T-DNA-induced mutations in other genes. By contrast, an independent T-DNA insertion mutant (*iri1-2*) in the annotated gene of *iri1-1* displayed a complete *iri* phenotype (Figs. 1B-C) and was concomitantly impaired in RBH-induced priming of callose defence (Fig. 1D). The *iri1-1* and *iri1-2* mutants carry a T-DNA insertion in the 5^th^ intron and the 2^nd^ intron of the *Lysine Histidine Transporter1* (*LHT1*; At5G40780; Fig. 1B; Supplemental Figs. 3A-B), respectively, which encodes a high-affinity amino acid transporter for acidic and neutral amino acids in roots and mesophyll cells (Chen and Bush, 1997; Hirner et al., 2006; Svennerstam et al., 2007).

### *IRI1/LHT1* expression controls RBH uptake from the soil

Since IRI1/LHT1 acts as an amino acid transporter (Chen and Bush, 1997), we hypothesised that the lack of RBH-IR in *iri1* is caused by impaired RBH uptake from the soil. To examine this hypothesis, we determined RBH leaf concentrations after soil-drench treatment with increasing RBH concentrations in Col-0, *iri1-1* and a previously characterised over-expression line of *IRI1/LHT1* (Hirner et al., 2006; *Pro35S:IRI1/LHT1*), which shows a 27-fold higher *IRI1/LHT1* expression level than Col-0 plants under our experimental conditions (Supplemental Fig. 3C). At 2 days after soil treatment, replicate leaf tissues were sampled for RBH quantification by hydrophilic interaction liquid chromatography coupled to quadrupole time-of-flight mass spectrometry (HILIC-Q-TOF; Fig. 2A) or challenged the leaves with *Hpa* to quantify RBH-IR (Fig. 2B). The three genotypes differed statistically in RBH shoot concentrations after soil treatment with increasing RBH concentrations, as evidenced by a highly significant statically interaction between soil treatment and genotype (2-way ANOVA; *p*<0.001; Fig. 2A). For both Col-0 and *Pro35S:IRI1/LHT1*, RBH shoot accumulation showed a dose-dependent rise with increasing RBH concentrations applied to the soil. The *Pro35S:IRI1/LHT1* plants accumulated statistically higher RBH concentrations than Col-0 at 0.15 and 0.5 mM RBH applied to the soil, whereas RBH concentrations in the shoot of *iri1-1* were hardly detectable by HILIC-Q-TOF and failed to show a dose-dependent increase by RBH soil treatment (Fig. 2A). The observed variation in RBH shoot concentrations correlated with the intensity of RBH-IR against *Hpa* (Fig. 2B); while RBH failed to induce statistically significant levels of resistance in *iri1-1* at all concentration tested, *Pro35S:IRI1/LHT1* plants showed increased levels of RBH-IR compared to Col-0 at all RBH concentrations tested. Notably, the relatively low concentration of 0.05 mM RBH failed to protect against *Hpa* in Col-0 plants, whereas the same RBH concentration in *Pro35S:IRI1/LHT1* resulted in a statistically significant reduction in *Hpa* colonisation (Fig. 2B). Thus, the IRI1/LHT1 protein seems solely responsible for the uptake of RBH from the soil and subsequent accumulation of this chemical in the shoot. Accordingly, the expression of the *IRI1/LHT1* gene determines the level of RBH-IR against *Hpa*.

**Figure 2.**
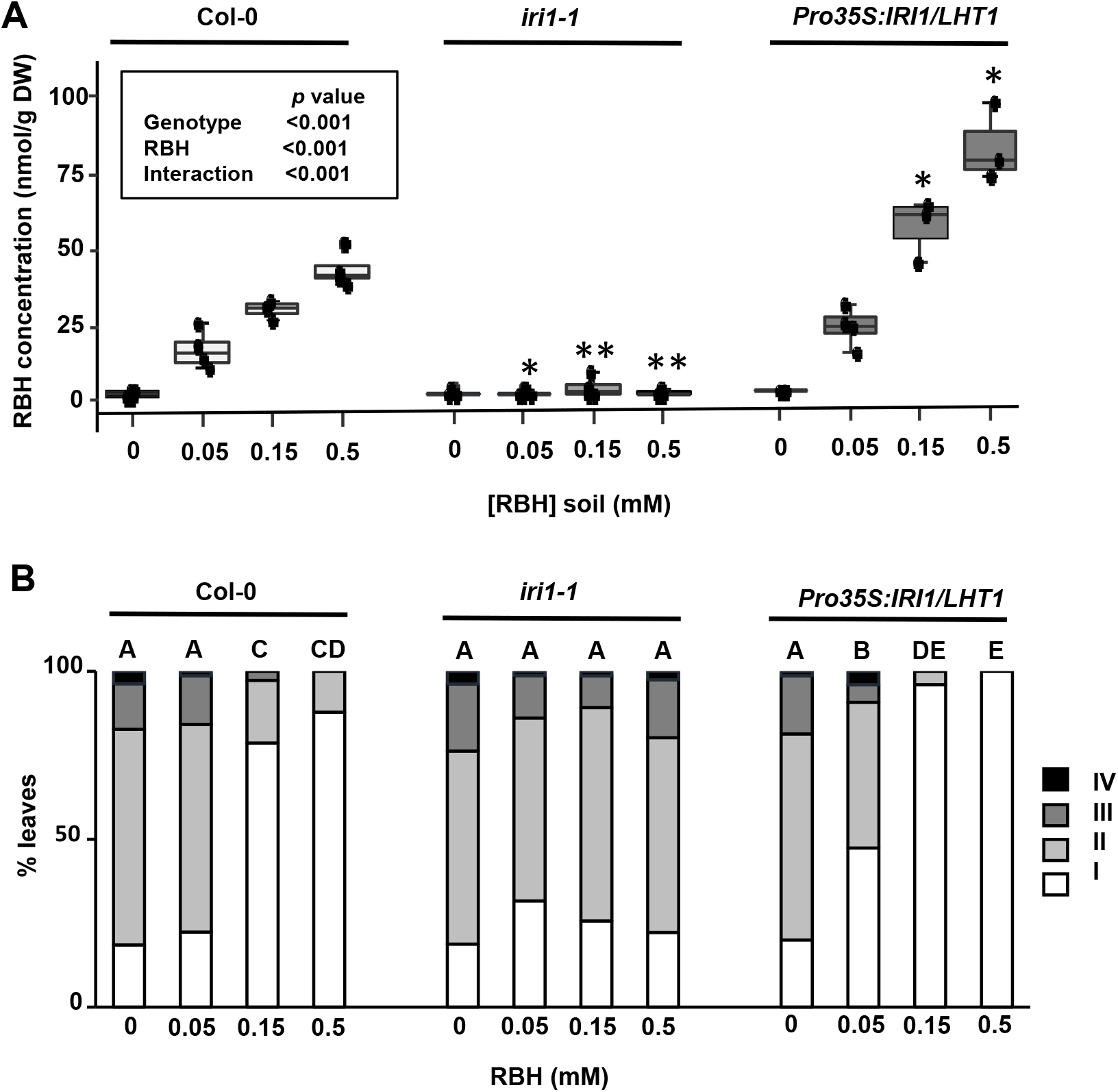
*IRI1/LHT1* controls RBH-uptake and RBH-induced resistance against *Hpa*. **(A)** Quantification of RBH in leaves of Col-0 (wild-type), *iri1-1/lht1-5* (mutant) and *Pro35S:IRI1/LHT1* (over-expression) plants after soil-drench treatment with increasing RBH concentrations. Leaves were collected at 2 days after soil-drench treatment with RBH and analysed by HILIC-Q-TOF. Boxplots show median (middle bar), interquartile range (IQR; box), 1.5 x IQR (whiskers) and replication units (single dots) of leaf RBH concentrations (nmol/g DW). Inset shows *p*-values of statistically significant effects on RBH concentration by genotype, soil treatment and interaction thereof (2-way ANOVA). Asterisks indicate statistically significant differences to Col-0 for each soil treatment (Welch t-test; **: *p*<0.001; *: 0.001<*p*<0.05). **(B)** Quantification of RBH-induced resistance against *Hpa* Col-0, *iri1-1/lht1-5* and *Pro35S:IRI1/LHT1*. Two-week-old seedlings were soil-drenched with increasing concentrations of RBH and challenge-inoculated with *Hpa* conidiospores 2 days later. Shown are frequency distributions of trypan-blue-stained leaves across four *Hpa* colonisation classes at 6 dpi (see Fig. 1A). Different letters indicate statistically significant differences between samples (Fisher’s exact tests + Bonferroni FDR; *p* < 0.05; n = 70-90 leaves).

### *IRI1/LHT* expression rather than catabolism controls plant tolerance to RBH

In contrast to BABA, RBH induces resistance in Arabidopsis without concomitant growth inhibition (Buswell et al., 2018). To examine whether *IRI1/LHT1* expression and RBH-uptake influence the tolerance of Arabidopsis to RBH, we quantified seedling growth of Col-0, *iri1-1*, and *Pro35S:IRI1/LHT1* on MS agar medium containing increasing concentrations of RBH in the presence of increasing concentrations of L-Alanine, a canonical substrate of IRI1/LHT (Hirner et al., 2006). As shown in Fig. 3, green leaf area (GLA) of Col-0 and *iri1-1* was unaffected by increasing concentrations of RBH after 1 week of growth, irrespective of the concentration of co-supplied L-Alanine (L-Ala). By contrast, *Pro35S:IRI1/LHT1* plants displayed dose-dependent growth repression with increasing RBH concentrations, which was antagonised by increasing concentrations of L-Ala supplied to the agar medium. These results suggest that the natural tolerance of Arabidopsis to RBH reported previously (Buswell et al., 2018) is determined by the level of *IRI1/LHT1* expression and RBH uptake. To confirm this notion, we repeated the experiment on MS medium without inorganic nitrogen (N_inorg_; NO_3_^-^ and NH_4_^+^), supplemented with increasing concentrations of RBH and L-Ala. While Arabidopsis failed to grow on agar medium without N_inorg_ (Supplemental Fig. 4), increasing RBH concentrations in the growth medium failed to rescue growth. Hence, Arabidopsis cannot metabolise RBH as a N source, which rules out metabolic breakdown (catabolism) as a mechanism of RBH tolerance. By contrast, increasing L-Ala concentrations applied to the agar medium rescued seedlings growth of all genotypes tested, albeit to varying degrees. While *Pro35S:IRI1/LHT1* plants showed the strongest growth response to increasing L-Ala concentrations, Col-0 displayed an intermediate growth response, followed by a relatively weak growth response in *iri1-1* (Supplemental Fig. 4), confirming the contribution of IRI1/LHT1 to L-Ala uptake (Hirner et al., 2006; Svennerstam et al., 2007; Svennerstam et al., 2011). Notably, increasing RBH concentrations in the presence of L-Ala caused dose-dependent growth reduction in *Pro35S:IRI1/LHT1* plants, whereas Col-0 and *iri1-1* did not display the same trend (Supplemental Fig. 4). This supports our conclusion that the enhanced expression of *IRI1/LHT1* in *Pro35S:IRI1/LHT1* renders the plant sensitive to RBH-induced stress, resulting in accumulation of phytotoxic RBH concentrations that cannot be metabolised. Hence, Arabidopsis tolerance to RBH is controlled by *IRI1/LHT1* gene expression rather than RBH catabolism.

**Figure 3.**
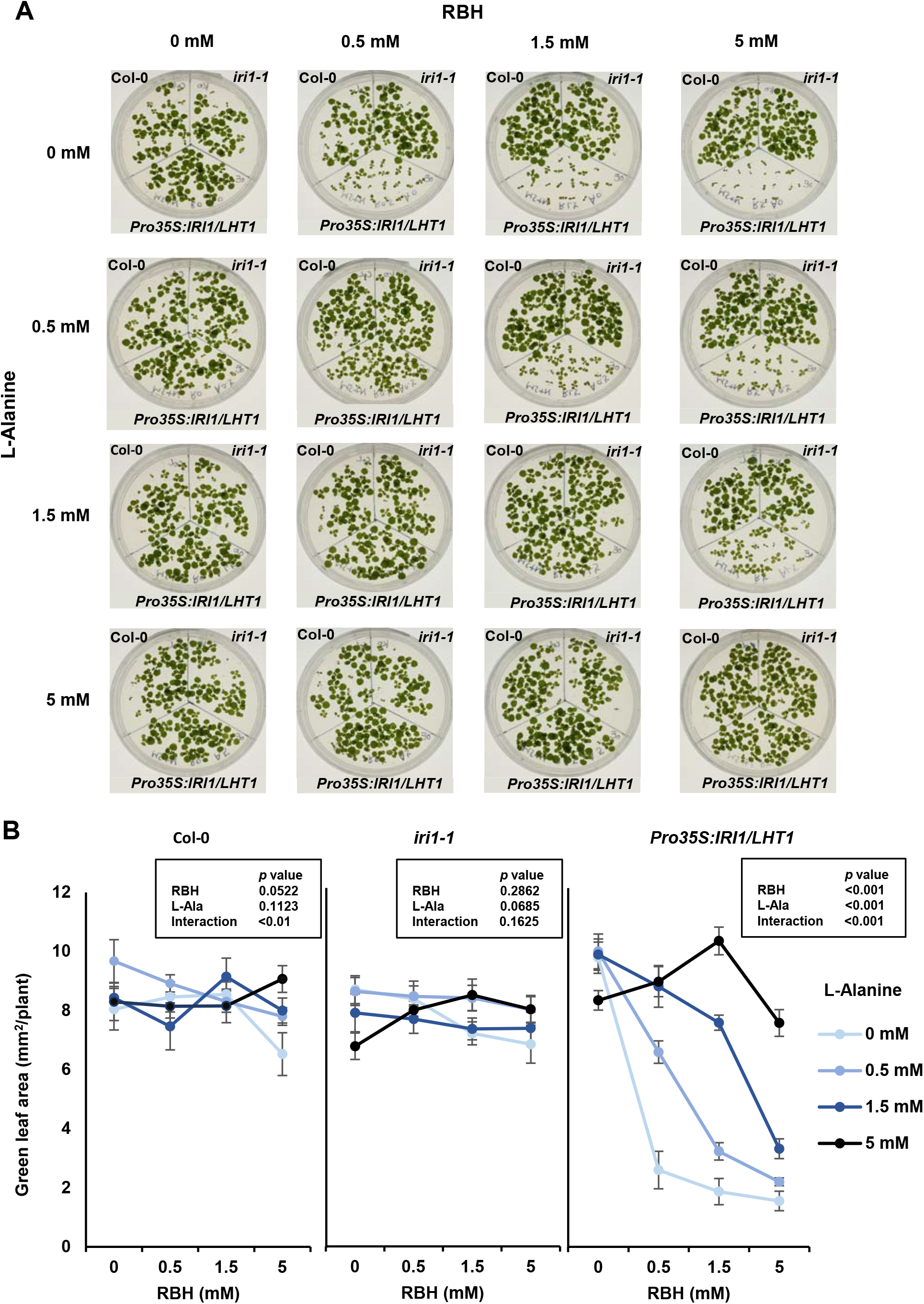
Over-expression of *IRI1/LHT1* renders Arabidopsis susceptible to growth repression by RBH, which is antagonised by co-application of L-Alanine. **(A)** *IRI1/LHT1*-dependent effects of RBH and L-Alanine on plant growth. Shown are 2-week-old seedlings of Col-0 (upper left), *iri1-1/iht1-5* (upper right), and *Pro35S:IRI1/LHT1* (bottom) grown on MS agar plates, supplemented with 10 mM (NH_4_)_2_SO_4_ and increasing concentrations of RBH and/or L-Alanine. **(B)** Quantification of green leaf area (GLA ± SEM; n=7-19) in 1-week-old Col-0, *iri1/iht1-5*, and *Pro35S:IRI1/LHT1* seedlings from the same experiment. Inset shows p-values of effects on green leaf area by RBH concentration, L-Alanine concentration and interaction thereof inside each genotype (2-way ANOVA).

### *IRI1/LHT1* expression controls BABA uptake, BABA-IR and BABA tolerance

Given the broad substrate range of the IRI1/LHT1 transporter for acidic and neutral amino acids (Hirner et al., 2006; Svennerstam et al., 2007), we examined whether IRI1/LHT1 also plays a role in the uptake of BABA. To this end, replicate shoot tissues of Col-0 and *iri1-1* were sampled for *in planta* quantification of BABA concentrations at 2 days after soil drenching with increasing BABA concentrations (0, 0.025, 0.05, 0.15 and 0.5 mM), using HILIC-Q-TOF (Fig. 4A). While soil-drench treatment of Col-0 with increasing BABA concentrations resulted in a dose-dependent increase of the BABA concentrations in the shoot (Figs. 4A), similar treatment of the *iri1-1* mutant failed to increase shoot BABA concentrations (Fig. 4A), indicating that BABA uptake is dependent on IRI1/LHT1. To further corroborate this, Col-0, *iri1-1* and *Pro35S:IRI1/LHT1* were soil-drenched with increasing BABA concentrations and analysed for BABA-IR against *Hpa* (Fig. 4B). As reported previously, BABA was more efficient than RBH in protecting Col-0 against *Hpa* (Buswell et al., 2018), already protecting against *Hpa* colonisation at 0.0025 mM BABA and reaching maximum levels of protection at concentrations of 0.05 mM and higher (Fig. 4B). The *Pro35S:IRI1/LHT1* line showed even higher levels of protection against *Hpa* colonisation at 0.025 mM BABA in comparison to Col-0, indicating that it is sensitised to respond to BABA. By contrast, the *iri1-1* mutant was severely compromised in its effectiveness of BABA-IR, and only displayed weak levels of IR at soil concentrations of 0.25 mM and 0.5 mM (Fig. 4B). Thus, like RBH-IR, BABA-IR is determined by the expression level of *IRI1/LHT1*, which supports our notion that LHT1/IRI1 controls BABA-uptake from the soil. Finally, to determine whether IRI1/LHT1 controls tolerance to BABA-induced phytotoxicity, we quantified growth of Col-0, *iri1-1* and *Pro35S:IRI1/LHT1* on agar plates supplemented with phytotoxic concentrations of BABA. As shown in Fig 5, GLA values of Col-0 after 1 week of growth declined with increasing BABA concentrations. This BABA-induced stress response was dramatically increased in *Pro35S:IRI1/LHT1* plants and reduced, although not abolished, in *iri1-1* plants (Fig. 5). The fact that *iri1-1* plants still showed growth repression at higher BABA concentrations suggests that there are additional mechanisms contributing to BABA-induced phytotoxicity. Nonetheless, the BABA experiments collective indicate that IRI1/LHT1 is the dominant transporter for BABA uptake from the soil. As such, the expression of the *IRI1/LHT1* gene not only controls the strength of the BABA-IR response but also the tolerance to BABA-induced stress.

**Figure 4.**
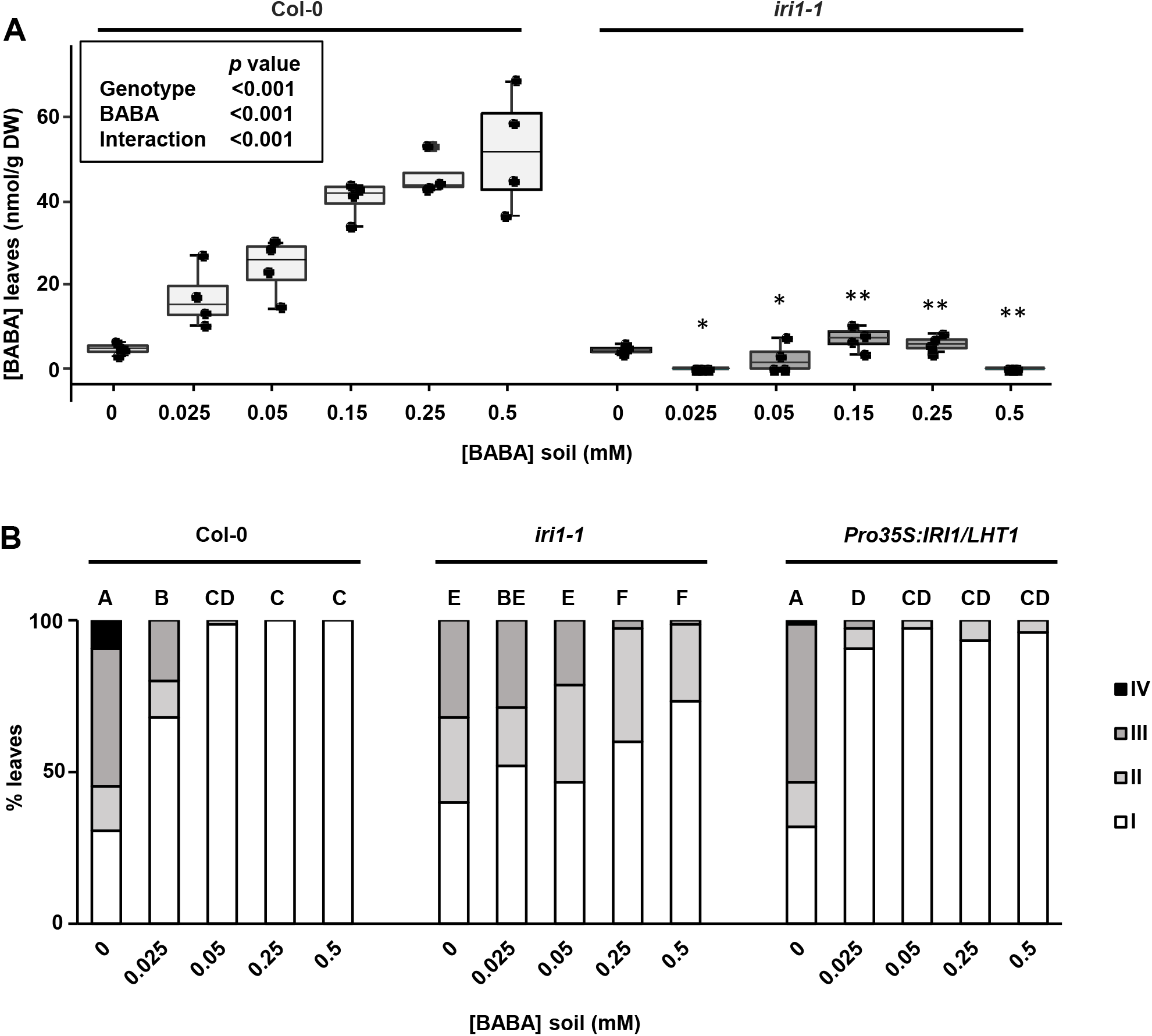
*IRI1/LHT1* controls BABA-uptake and BABA-induced resistance against *Hpa*. **(A)** Quantification of BABA in leaves of Col-0 (wild-type) and *iri1-1/lht1-5* (mutant) plants after to soil drench treatment with increasing BABA concentrations. Leaves were collected at 2 days after soil-drench treatment and analysed by HILIC-Q-TOF. Boxplots show median (middle bar), interquartile range (IQR; box), 1.5 x IQR (whiskers) and replication units (single dots) of leaf BABA concentrations (nmol/g DW). Inset shows *p*-values of statistically significant effects on BABA concentration by genotype, soil treatment and interaction thereof (2-way ANOVA). Asterisks indicate statistically significant differences to Col-0 for each soil treatment (Welch t-test; **: *p*<0.001; *: 0.001<*p*<0.05). **(B)** Quantification of BABA-induced resistance against *Hpa* in Col-0, *iri1-1/lht1-5* and *Pro35S:IRI1/LHT1* plants. Two-week-old seedlings were soil-drenched with increasing concentrations of BABA and challenge-inoculated with *Hpa* conidiospores 2 days later. Shown are frequency distributions of trypan-blue-stained leaves across four *Hpa* colonisation classes at 6 dpi (see Fig. **1A**). Different letters indicate statistically significant differences between samples (Fisher’s exact tests + Bonferroni FDR; *p* < 0.05; n = 70-80 leaves).

**Figure 5.**
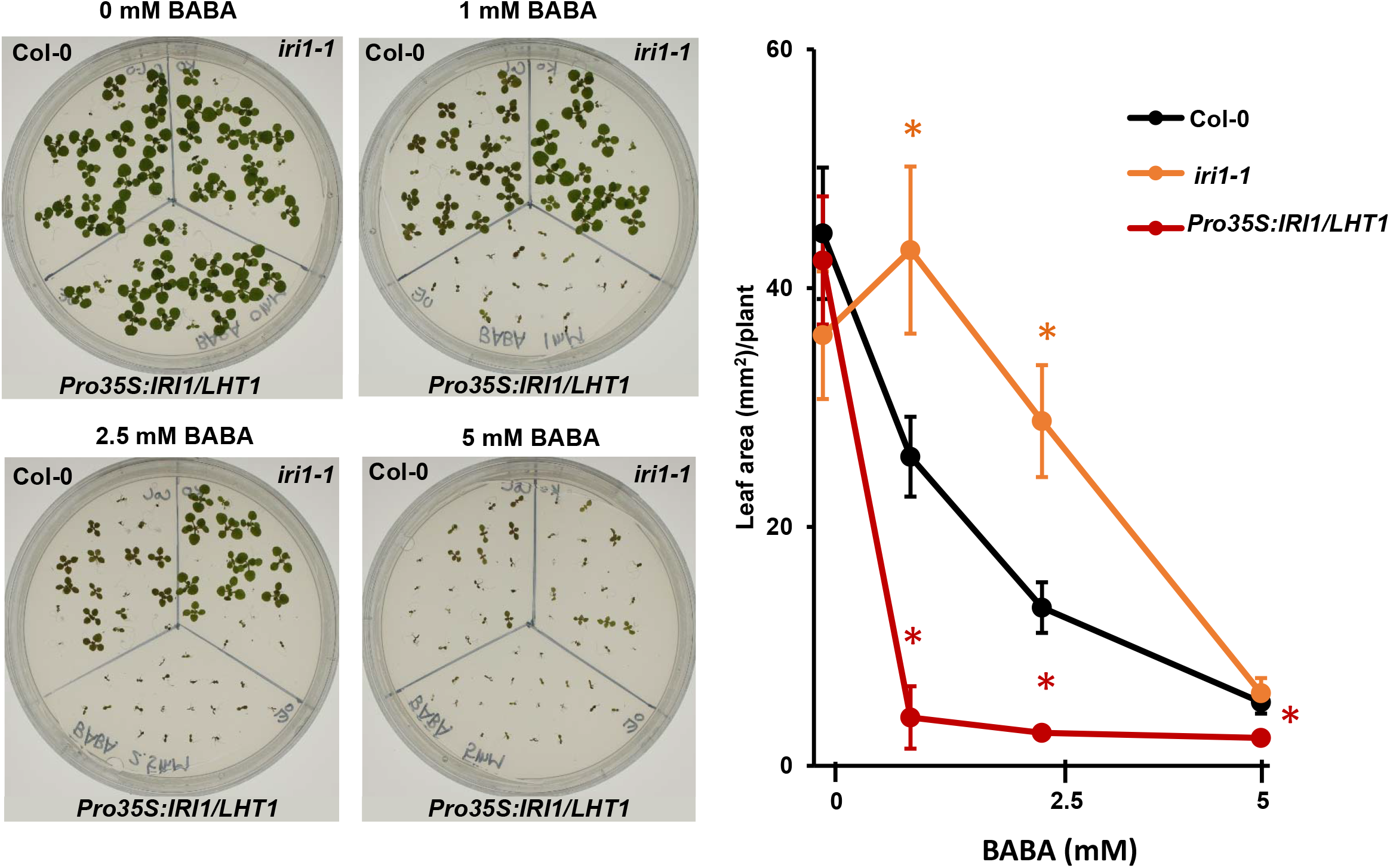
*IRI1/LHT1* controls stress tolerance to BABA. **(A)** Effects of BABA on growth by Col-0, *iri1-1/lht1-5, Pro35S:IRI1/LHT1* Shown are 2-week-old seedlings of Col-0 (upper left), *iri1-1/iht1-5* (upper right), and *Pro35S:IRI1/LHT1* (bottom) grown on MS agar plates, supplemented with increasing concentrations of BABA. **(B)** Average green leaf areas (GLA ± SEM; n=9-20) of 1-week-old Col-0, *iri1-1/lht1-5, Pro35S:IRI1/LHT1* plants from the same experiment. Asterisks indicate statistically significant differences compared to Col-0 at each BABA concentration (Welch t-tests + Bonferroni FDR; *p* < 0.05).

### LHT1 is a high-affînity transporter of both RBH and BABA

Having established that IRI1/LHT1 is responsible for the uptake of RBH and BABA, we next examined the kinetics by which IRI1/LHT1 transports these β-amino acids. To this end, the Arabidopsis *IRI1/LHT1* gene was heterologously expressed in the 22Δ10α strain of yeast, a mutant that is completely deficient in the uptake of amino acids (Besnard et al., 2016). In contrast to empty vector (EV)-transformed 22Δ10α cells, the *IRI1/LHT1*-expressing 22Δ10α strain was capable of growing on agar plates containing 1 mM L-Ala as the only nitrogen (N) source (Fig. 6A), while supplementing liquid growth medium without inorganic (NH_4_)_2_SO_4_ with increasing L-Ala concentrations steadily improved planktonic growth by *IRI1/LHT1*-expressing 22Δ10α cells (Fig. 6B). Increasing RBH and BABA concentrations in liquid growth medium with 1 mM L-Ala repressed growth by *IRI1/LHT1*-expressing 22Δ10α cells completely (Fig. 6C and D), despite the fact that both chemicals only marginally repressed 22Δ10α growth in liquid medium with 10 mM (NH_4_)_2_SO_4_ as N source (Supplemental Fig. 5). These results not only show that yeast fails to metabolise RBH and BABA, but they also suggest that increasing RBH and BABA concentrations outcompete L-Ala for cellular uptake.

**Figure 6.**
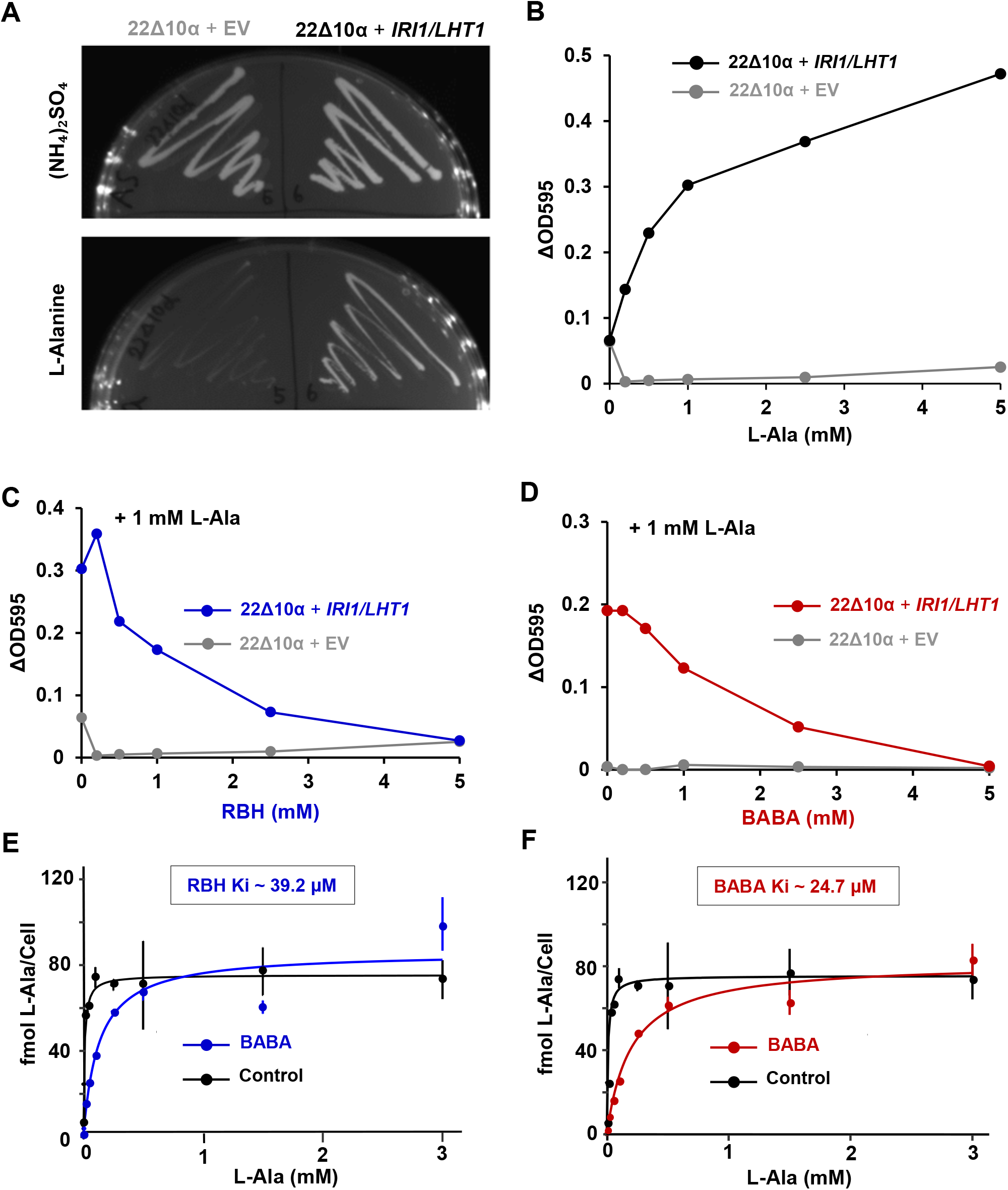
Characterisation of RBH- and BABA-uptake kinetics by IRI1/LHT1 via heterologous expression in yeast. **(A)** Transformation of the yeast mutant 22Δ10α (Besnard et al., 2016) with the *IRI1/LHT1* gene of Arabidopsis conplements growth on agar medium with L-Alanine (L-Ala) as the only N source. Shown are growth phenotypes of empty vector (EV)- and *IRI1/LHT1*-transformed 22Δ10α cells on agar medium supplemented with inorganic nitrogen (10 mM (NH_4_)_2_SO_4_; top) or 1 mM L-Alanine (bottom). **(B - D)** Planktonic growth of EV- and *IRI1/LHT1*-transformed 22Δ10α in liquid medium supplemented with increasing L-Ala concentrations (**B**), 1 mM L-Ala + increasing RBH concentrations (**C**), or 1 mM L-Ala + increasing BABA concentrations (**D**). Growth was quantified spectrophotometrically by ΔOD_595_ values (OD_595_ - OD_595_ of medium without yeast). Shown are average ΔOD_595_ values ±SEM (n=4) after 3 days of growth. **(E - F)** Competitive inhibition of IRI1/LHT1-dependent uptake of L-Ala by RBH (E; blue) and BABA (F; red). Uptake velocities by IRI1/LHT1 were determined in the presence of increasing L-Ala concentrations containing 50 nCi ^14^C-labelled L-Ala with and without 500 μM RBH (**E**) or BABA (**F**). Data represent average L-Ala uptake velocities (fmol L-Ala/cell ±SEM; n=3) over a 5 min. time-window. In the absence of RBH or BABA, the Km for L-Ala-uptake by IRI1/LHT1 was 9.4 μM. Competitive inhibition by RBH and BABA is shown by a decrease in Km but not V_max_. Insets indicate inhibitor constants (Ki) for RBH (39.3 μM) and BABA (24.7 μM).

To study the kinetics of RBH and BABA uptake, we carried out experiments with ^14^C-labelled L-Ala in the absence and presence of RBH or BABA. To this end, EV- and *IRI1/LHT1*-expressing 22Δ10α cells were incubated for 2, 5 and 10 min. in buffer containing 50 or 500 μM L-Ala with a fixed amount of ^14^C-L-Ala of incubation, after which cellular L-Ala uptake was quantified by ^14^C scintillation. In contrast to EV-transformed cells, IRI1/LHT1-expressing cells showed linear uptake of L-Ala over time (Supplemental Fig. 6), confirming the functionality of the transporter in yeast. To determine whether RBH and BABA competitively inhibit the IRI1/LHT1 transporter for L-Ala uptake, IRI1/LHT1- expressing cells were incubated for 5 min. in buffer containing increasing concentrations L-Ala with a fixed amount of ^14^C-L-Ala in the presence and absence of 500 μM RBH or 500 μM BABA (Fig. 6E and F). Plotting the uptake velocity (fmol L-Ala/cell) against L-Ala concentration revealed a dose-dependent increase until saturation (V_max_; Fig. 6E and F). Based on these data, we calculated that IRI1/LHT1 has a Km value of 9.4 μM for L-Ala-uptake, which is in line with previously reported Km values for acidic and neutral amino acids (Hirner et al., 2006). Although uptake velocity in the presence of either 500 μM RBH or 500 μM BABA was repressed across a lower range L-Ala concentrations, it still reached maximum levels at higher L-Ala concentrations, indicating that RBH and BABA are competitive inhibitors of L-Ala uptake by IRI1/LHT1. Based on these data, we calculated an inhibition constant (Ki) of 39.2 μM and 24.7 μM for RBH and BABA, respectively, which is within similar range as Km values of other α-amino acids (Hirner et al., 2006). Hence, IRI1/LHT1 is a high-affinity transporter of both RBH and BABA, with a higher affinity for BABA than for RBH.

## DISCUSSION

### Using annotated T-DNA insertion lines for a genome-saturating mutant screen

We have used a genome-covering collection of Arabidopsis T-DNA insertions lines in a forward mutant screen for regulatory genes of IR. The availability of homozogous T-DNA insertions with high genomic coverage (Alonso and Ecker, 2006) ensures a near genome-saturating screen. The use of this resource has several benefits compared to conventional mutant screens. Firstly, the availability of T-DNA flanking sequences mapped to the Arabidopsis genome allows for immediate identification of gene candidates without having to commit to time-consuming generation of mapping populations and linkage analysis. Secondly, the collection of homozygous mutant lines enables screening of small populations that all carry the same mutant allele, which facilitates the identification of partial (leaky) mutant phenotypes as illustrated by the selection of 104 *iri* lines that were are partially affected in RBH-IR (Fig. 1A; Supplemental Fig. 1, Supplemental Data Set 1). This relatively high number of partial *iri* mutants supports the notion that IR is a highly quantitative form resistance, relying on the additive contribution of multiple genes (Ton et al. 2006; Ahmad et al. 2010, Wilkinson et al. 2019). Thus, the within-genotype replication of this screen enables selection for genes that make a quantitative contribution to complex multigenic traits. A disadvantage of using annotated T-DNA insertion lines in a forward mutant screen is that a single T-DNA insertion line can carry multiple mutations (O’Malley et al., 2015). These mutations are not necessarily covered by the annotated T-DNA flanking sequences, since they can be caused by truncated T-DNA elements or mis-repairs of integration sites from abortive T-DNA integrations (‘mutational foot prints’; Gelvin, 2021). Indeed, several other studies have reported that mutant phenotypes in this collection of T-DNA insertion lines do not always co-segregate with the annotated T-DNA insertion (De Muyt et al., 2009; Dobritsa et al., 2011; Wilson-Sánchez et al., 2014). To account for this, we validated all the mutant phenotypes of the 4 complete *iri* mutants in independent T-DNA insertion lines of annotated genes for both RBH-IR and augmented cell wall defence against *Hpa* (Fig. 1C and Supplemental Fig. 2). Even though the *iri* phenotypes of the 4 original mutant lines were robust and reproducible (Fig. 1C and Supplemental Fig. 2), only the phenotype of the *iri1-1* mutant could be confirmed in an independent T-DNA insertion line in the annotated gene. Identifying the causal mutation in the other 3 *iri* lines would requires further TAIL-PCR to identify flanking sequences of alternative T-DNA insertions, or conventional linkage analysis in segregating mapping populations.

### The role of LHT1/IRI1 in plant-biotic interactions

The *IRI1* gene encodes the broad-range amino acid transporter LHT1. Cellular transporters play important roles in the control of plant-pathogen interactions by facilitating pathogen feeding (Elashry et al., 2013; Marella et al., 2013), secretion of antibiotic compounds (Lu et al., 2015; Khare et al., 2017), transporting defence hormones (Serrano et al., 2013), or contributing to plant defence responses (Liu et al., 2010; Yang et al., 2014). Furthermore, the *IRI1/LHT1* ortholog *LjLHT1.2* of *Lotus japonicus* is transcriptionally induced by by arbuscular mycorrhizal fungi (AMF; Guether et al., 2011), suggesting it facilitates AMF-dependent uptake of organic nitrogen. Given the role of IRI1/LHT1 in IR, it is tempting to speculate that LHT1 also plays a role in mycorrhiza-IR (Cameron et al., 2013). In Arabidopsis. *IRI1/LHT1* has been implicated directly in the regulation of SA-dependent disease resistance. Liu et al. (2010) reported that *iri1/lht1* mutant lines have increased basal resistance against the hemibiotrophic bacterium *Pseudomonas syringae* pv. *tomato*, the hemibiotrophic fungus *Colletotrichum higginsianum* and the biotrophic fungus *Erysiphe cichoracearum*. The study furthermore provided evidence that *IRI1/LHT1* controls plant immunity by cellular uptake of L-Glutamine (L-Gln), which is a precursor of the redox-buffering compound glutathione. Liu et al. (2010) proposed that the reduced L-Gln uptake capacity in *iri1/lht1* mutants reduces redox buffering capacity, thereby enabling augmented elicitation of ROS and SA-dependent defences upon pathogen attack. However, our experiments did not revealed statistically significant differences in basal defence against the biotrophic Oomycete *Hpa* between wild-type and *iri1/lht1* mutant plants (Fig. 1 and Supplemental Fig. 2). This discrepancy could be explained by the fact that we used relatively young plants (2- to 3-week-old seedlings), which do not express SA-dependent age-related resistance (ARR; Kus et al., 2002). Accordingly, it is possible that glutamine-dependent redox regulation contributes to age-related resistance in older plants. Since *IRI1/LHT1* expression is lower in younger plants (Hirner et al., 2006), it is also possible that there are other amino transporters contributing to the cellular delivery of glutamine in younger seedlings, such as AAP1 (Boorer et al., 1996) or CAT8 (Yang et al., 2010). Interestingly, in contrast to the negative role of *IRI1/LHT1* in innate immunity reported by Liu et al. (2010), a recent study by Yoo et al. (2020) revealed that *IRI1/LHT1* contributes positively to ETI-related resistance in Arabidopsis against *Pseudomonas syringae* pv. *maculicola* carrying the avirulence gene *AvrRpt2*. It should be noted, however, that the immune-related function of *IRI1/LHT1* described in our study is related to IR by the priming-inducing β-amino acids, rather than age-related basal resistance or ETI.

### The role of LHT1/IRI1 in beta-amino acid-IR

Our results have shown that *IRI1/LHT1* is the dominant transporter for cellular uptake of RBH and BABA from the soil (Figs. 2 and 4). LHT1/IRI1 is localised to the cell membrane (Hirner et al., 2006), which enables cellular import of RBH and BABA from the apoplast. The *IRI1/LHT1* gene is expressed in root tips, lateral roots and mature leaves (Hirner et al., 2006), enabling cellular uptake of RBH and BABA in both roots and leaves. Since *IRI1/LHT1* is not expressed in the leaf vein, we propose that the activity of RBH and BABA in leaves is preceded by long-distance transportation via the xylem and apoplastic distribution in the leaves. While BABA was applied exogenously in our experiments, recent studies have reported that biotic and abiotic stresses can elicit low concentrations of endogenous BABA in Arabidopsis (Thevenet et al., 2017; Balmer et al., 2019). Under these conditions, BABA only accumulates in locally stressed tissues, and not in systemic non-stressed tissues (Balmer et al., 2019), indicating that stress-induced accumulation of BABA does not contribute to systemic defence signalling. Although the biosynthesis pathway of stress-induced BABA remains unknown, it seems plausible that this local biosynthesis occurs inside the cell. This in turn suggests that cellular uptake of BABA by IRI1/LHT1 does not play a major role in the activity of stress-induced BABA, which would also explain why the *iri1* mutant and *Pro35S:IRI1/LHT1* over-expression lines in our experiments were not majorly affected in basal resistance to *Hpa* (Fig. 1 and Supplemental Fig. 2).

While our results provide strong evidence that IRI1/LHT1 is the uptake transporter of RBH and BABA (Figs. 2-6), this does not necessarily mean that the contribution of IRI1/LHT1 to RBH-/BABA-IR solely depends on uptake. For instance, while treatment with 0.05 mM RBH resulted in similar foliar concentrations in both *Pro35S:IRI1/LHT1* and wild-type plants (Fig. 2A), this relatively low RBH concentration only triggered a significant IR response in *Pro35S:IRI1/LHT1* plants and not in wild-type plants. This uncoupling of RBH concentration from IR suggests that the function of IRI1/LHT1 in RBH-IR may involve an additional defence signalling activity that becomes active at low RBH concentrations. Such a transporter-receptor co-functionality (‘transceptor’ activity) has been reported for NRT1.1 for nitrate uptake and signalling. Replacing Pro492 with Leu492 in NRT1.1 disabled the nitrate transporting activity of this protein but not its ability to induce *NRT2.1* expression (Ho et al., 2009), which is a nitrate-responsive gene that has concomitantly been linked to the regulation of disease resistance (Camanes et al., 2012). Although no amino acid transporters have been reported with receptor co-functionality (Dinkeloo et al., 2018), it is tempting to speculate that IRI1/LHT1 acts as a transceptor of β-amino acids. Site-directed mutagenesis of IRI1/LHT1 and testing whether its RBH and BABA transportation activity can be uncoupled from its role in RBH-/BABA-IR would be required to validate this attractive hypothesis.

### RBH and BABA compete with proteinogenic amino acids for uptake by IRI1/LHT1

We have used *IRI1/LHT1*-expressing yeast cells to assess competitive inhibition of L-Ala uptake by RBH and BABA. Our uptake essays revealed a Km of IRI1/LHT1 of 9.4 μM for L-Ala (Fig. 6E), which is similar to previously reported Km values of IRI1/LHT1 for proteinogenic amino acids (Hirner et al., 2006). Furthermore, the inhibitory kinetics of RBH or BABA on L-Ala uptake confirmed competitive inhibition, as evidenced by the fact that L-Ala uptake in the presence of RBH or BABA still reached maximum velocities at higher L-Ala concentrations (Fig. 6E-F). Of the two beta-amino acids, BABA had a lower Ki than RBH (24.7 μM vs 39.2 μM), suggesting IRI1/LHT1 has a higher affinity for BABA than for RBH (Figs. 6E-F). As IRI1/LHT1 has been shown to have a similar affinity for a range of acidic and neutral amino acids, including L-Gln (Hirner et al., 2006; Svennerstam et al., 2007), our results also explain previous findings by Wu et al. (2010), who reported that BABA-induced phytotoxicity in Arabidopsis can be alleviated by co-application with L-Gln.

### LHT1/IRI1: not just a transporter for proteinogenic amino acids

Although IRI1/LHT1 was initially identified as a transporter for proteinogenic amino acids (Chen and Bush, 1997), subsequent studies have shown that it transports a much wider range for non-proteinogenic amino acids, such as the ethylene precursor 1-aminocyclopropane-1-carboxylate (ACC; Shin et al., 2015) and xenobiotic amino acid conjugates (Chen et al., 2018; Jiang et al., 2018). Consistent with this broad-spectrum uptake activity, we have shown that IRI1/LHT1 is the main transporter of the β-amino acids RBH and BABA. Of particular interest is the regulatory function of IRI1/LHT1 in the trade-off between beta-amino acid-IR and plant growth. While over-expression of *IRI1/LHT1* in *Pro35S:IRI1/LHT1* plants increased BABA-IR at the relatively low concentration of 0.025 mM BABA (Fig. 4), it also dramatically increased the plant’s sensitivity to BABA-induced growth repression (Fig. 5). By contrast, RBH elicited high levels of IR at soil concentrations of 0.15 mM RBH and above (Fig. 2B), but did not repress growth across a range of concentrations (Fig. 3), which supports our earlier conclusion that RBH induces disease resistance without costs on plant growth (Buswell et al. 2018). Interestingly, however, over-expression of *IRI1/LHT1* gene in *Pro35S:IRI1/LHT1* plants increased levels of IR at relatively low RBH concentrations (Fig. 2B), but also repressed growth in a dose-dependent manner (Fig. 3). These findings reveal two important conclusions. Firstly, like BABA, RBH is able to repress plant growth, but this phytotoxicity depends of the plant’s uptake capacity, which in turn is determined by the level of *IRI1/LHT1* gene expression (Supplemental Fig. 3C). This conclusion is consistent with our other finding that Arabidopsis fails to metabolise RBH as an N source (Supplemental Fig. 4). Secondly, our results show that the trade-off between plant growth and beta-amino acid-IR can be optimised in favour of the IR response by manipulating the level of *IRI1/LHT1* gene expression. This conclusion holds major translational value for breeding programmes aiming to exploit BABA-IR in vegetable crops that are protected by BABA but also suffer from BABA-induced phytotoxicity (Cohen et al., 2016; Yassin et al., 2021).

## METHODS

### Biological material

All *Arabidopsis thaliana* genotypes were in the genetic background of accession Col-0. The *iri1-1* mutant (*lht1-5*) and *iri1-2* mutant (*lht1-4*) was described previously by Liu et al. (2010); the *Pro35S:IRI1/LHT1* overexpression lines was described by Hirner et al. (2006). The *iri* mutant screen was performed with fully annotated homozygous T-DNA insertion lines from the SALK and SAIL collections (Alonso *et al*., 2003) and purchased from the Nottingham Arabidopsis Stock Centre (sets N27941, N27951, N27942, N27943, N27944, N27945). The annotated T-DNA insertions in *iri1-1* (SALK_115555)*, iri1-2* (SALK_036871)*, iri2-1* (SALK_204380), SAIL_902_B08, *iri3-1* (SALK_118654), SALK_078838*, iri4-1* (SALK_076708) and SALK_046376 were confirmed by PCR before further testing (Supplemental Table 1), as described below. *Hyaloperonospora arabidopsidis* strain WACO9 was maintained in its asexual cycle by alternate conidiospores inoculations of Col-0 and Ws *NahG* plants.

### Plant growth conditions

For soil-based IR experiments, seeds were planted in a 2:1 (v/v) Scott’s Levington M3 compost/sand mixture and stratified for 2-4 days in the dark at 4 °C. Plants were subsequently cultivated under short-day conditions (8 h-day; 150 μmol photons m^-2^ s^-1^; 21 °C; and 16 h-night; 18 °C) at ~60% relative humidity (RH). Plants for seed propagation were grown at long-day growth conditions (16 h-day; 150 μmol photons m^-2^ s^-1^; 21 °C; and 8 h-night; 18 °C) at ~60% RH. For plate assays, seeds were surface-sterilised (vapour-phase sterilisation) prior to planting on ½-strength Murashige & Skoog (MS) 1.5% agar plates.

### Mutant screen

Approximately 10-15 seeds of each seed line were planted in individual wells of 400-well trays (Teku JP 3050/230 H). Each tray was filled with ~2.4 L of compost/sand mixture. After planting, stratification of seeds and seed germination, seedling densities were equalised to 5 seedlings/well. Two-week-old seedlings were treated with RBH by adding 1,5 L of 2x concentrated RBH solution (1mM) to each tray, which was left overnight to saturate the soil. Excess RBH solution (~300 mL) was removed the next morning, resulting in a final soil concentration of ~0.5mM RBH. Challenge-inoculation was performed at 2 days after RBH treatment by spraying seedlings with a suspension of *Hpa* conidiospores (10^5^ spores/mL). Trays were sealed with clingfilm after inoculation to maintain 100% RH and promote infection. To verify RBH-IR, each tray contained 3 randomly distributed wells with Col-0 seedlings. Furthermore, to verify favourable conditions for *Hpa* disease, 3 additional wells with Col-0 seedlings were cut from each tray and left outside during RBH-uptake to prevent RBH-IR prior to inoculation. At 5-7 dpi, trays were visually inspected for *Hpa* sporulation when sporulation on Col-0 seedlings in the untreated wells of the tray became apparent. Lines developing sporulation within 7 dpi were logged as stage 1 *<u>i</u>mpaired in <u>R</u>BH-induced-<u>i</u>mmunity* (S1 *iri*) lines, while un-germinated lines were logged as stage 1 ungerminated (S1 *ug*). All S1 *iri* and S1 *ug* lines were pooled for the stage 2 screen in 400-well trays, as described above. S1 *iri* lines allowing visible sporulation in two screens time were logged as Stage 2 *iri* (S2 *iri*). S1 *ug* lines that germinated upon rescreening and showing sporulation were re-tested for S2 *iri* phenotypes. Of the 26,631 T-DNA insertion lines, 23,547 lines germinated and could be screened for *iri* mutant phenotypes. The 427 putative *iri1* lines selected after stage 2 were pooled for seed bulking and validated by controlled IR assays in stage 3 (S3) of the screen, as described below.

### Induced resistance (IR) assays

Two-week-old plants were grown in 60-mL pots and soil-drenched with water, (*R*)-β-homoserine (Sigma-Aldrich; #03694), or R/S-BABA (Sigma-Aldrich, #A44207) to the indicated soil concentrations as described previously (Buswell et al., 2018). Two days after chemical treatment, plants were spray-inoculated with a suspension of *Hpa* conidiospores (10^5^ spores/mL) and maintained at 100% RH to promote infection. Leaves were collected at 6-7 dpi for trypan blue staining for microscopic scoring of *Hpa* colonisation by categorising them into 4 classes, ranging from healthy leaves (I) to heavily colonised leaves (IV), as described in detail by Schwarzenbacher et al. (2020). To investigate augmented induction of cell wall defence by chemical priming treatment, leaves were harvested at 3 dpi for aniline blue/calcofluor staining and analysis by epi-fluorescence microscopy (Leica DM6B; light source: CoolLED pE-2; 365 nm excitation filter, L 425 nm emission filter, 400 nm dichroic filter). For each genotype/treatment combination, germinated coniodiospores on 10 leaves from independent plants were scored either as arrested (spores or germ tubes fully encased in callose), or non-arrested by callose depositions (no callose or lateral callose deposition along the germ tube/hyphae), as detailed by Schwarzenbacher *et al*. (2020). Statistical differences in in *Hpa* colonisation or callose defence were analysed by pairwise Fisher’s exact tests, using R software (v 3.5.1). For multiple comparisons, an additional Bonferroni multiple correction was applied, using the R package ‘fifer’ (fifer_1.1.tar.gz).

### Plant growth assays

Surface-sterilised seeds were planted on ½-strength MS agar plates and cultivated for 2 weeks under standard plant growth conditions, as indicated above. Photos were taken after 1 and 2 weeks of growth with Nikon D5300. Green leaf areas (GLA) were quantified from digital photos of 1 or 2-week-old plants, using Fiji/ImageJ software (Rueden et al., 2017). Statistical differences in the natural logarithm of 1+green leaf area were analysed by 2-way ANOVA, using R software (v 3.5.1).

### Genotype verification by PCR and gene expression analysis by real-time quantitative PCR

Genomic T-DNA insertions of all *iri1, iri2, iri3 and iri4* lines were confirmed by PCR using LP+RP and LBb1.3/ LB3+RP primers (Supplemental Table S2) To quantify *IRI1/LHT1* gene expression by qRT-PCR, shoot tissues from five 2-week-old plants were collected and combined as 1 biological replicate. A total of 5 biological replicates were collected and snap-frozen in N_2_ (l) and homogenised. Total RNA was extracted using the RNeasy Plant Mini Kit (Qiagen, cat. no. 74904) and cDNA was synthesised from 800 ng RNA using the Maxima First Strand cDNA Synthesis Kit (Thermo Fisher, cat. no. K1641). The cDNA was diluted 20 times in nuclease-free water before qPCR. All qPCR reactions were performed with primer concentrations at a final concentration of 250 nM in a Rotor-Gene Q real-time PCR cycler (Qiagen, Q-Rex v1.0), using the Rotor-Gene SYBR Green PCR Kit (Qiagen, cat. no. 204074). The qPCR amplification of *IRI1/LHT1* was performed with gene-specific primers (FP: ATCTCCGGCGTTTCTCTTGCTG, RP: GCCCATGCGATTGTTGAGTAGCTG) and normalised to the transcript values of two housekeeping genes (At1g13440, and At2g28390), as detailed previously (Schwarzenbacher et al., 2020).

### Quantification of *in planta* RBH and BABA concentrations by hydrophilic interaction liquid chromatography coupled to quadrupole time-of-flight mass spectrometry

Shoot tissues were collected at 2 days after soil-drenching and divided into 4 replicate tubes per treatment (5 plants per tube, divided from separate trays), frozen at −80 °C, freeze-dried and weighed. Dry tissue was crushed and extracted into 1 mL of cold extraction buffer (MeOH:H_2_O:Formic acid, 10:89.99:0.01, v/v/v). Extracts were centrifuged at 4°C (16,000 g, 5 min.), after which each supernatant was divided between 3 aliquots. RBH and BABA standards were prepared as individual standards in the range from 0.1 to 100 μM. Separation was performed with a Waters Acquity HILIC BEH C18 analytical column, 1.7 mm particle size, 2.1 x 50 mm. The mobile phase was 20 mM ammonium formate with 0.1% formic acid (A) and acetonitrile with 0.1% formic acid (B). The gradient started at 99% A and reached 65% A in 4 min. The gradient changed to 1% A up to 6 min and was held there for 1.5 min and then returned to initial conditions. The solvent flow rate was 0.3 mL min^-1^, with an injection volume of 4 μL. Mass spectra were recorded in positive electro-spray ionisation mode, using a Waters UPLC system interfaced to a Waters quadrupole time-of-flight mass spectrometer (Q-TOF; G2Si Synapt). Nitrogen was used as the drying and nebulising gas. Desolvation gas flow was adjusted to approximately 150 L/h and the cone gas flow was set to 20 L/h with a cone voltage of 5 V and a capillary voltage of 2.5 kV. The nitrogen desolvation temperature was 280 °C and the source temperature was 100 °C. The instrument was calibrated in 20-1,200 m/z range with a sodium formate solution. Leucine enkephalin (Sigma-Aldrich, St. Louis MO, USA) in methanol:water (50:50) with 0.1% Formic acid was simultaneously introduced into the qTOF instrument via the lock-spray needle for recalibrating the *m/z* axis. Quantification of amino acids in tissues was based on the standard curves, using MassLynx v4.1 software (Waters, Elstree UK). Amino acids identities were confirmed by co-elution of product fragment ions with parent ions and matching peak retention times to individual amino acid standards. Statistical differences in RBH and BABA between genotypes and soil-drench treatments were tested by 2-way ANOVA followed by Welch t-tests to test cross-genotype differences at each RBH/BABA concentration, using R software (v 3.5.1).

### Yeast transformation

The *IRI1/LHT1* (At5g40780) gene with stop codon was amplified from wildtype Col-0 cDNA by Phusion High-Fidelity DNA Polymerase (New England Biolabs, #M0530L) and cloned into the pENTR plasmid (Invitrogen). *IRI1/LHT1* was then subcloned into pDR196 (Meyer et al., 2006) by restriction (EcoRI and XhoI) and ligation (T4 DNA ligase). Empty vector (EV)- and *IRI1/LHT1*-trandormed plasmids were confirmed by Sanger sequencing) and introduced into competent cells of the 22Δ10α strain (Besnard et al., 2016), using heat shock transformation (Gietz and Schiestl, 2007).

### Yeast growth assays

To assess growth of *IRI1/LHT1-* and EV-transformed 22Δ10α yeast strains, cells were first cultivated in liquid Yeast Nitrogen Base medium (Alfa Aesar, #H26271, without amino acids and ammonium sulfate) supplemented with 10 mM ammonium sulfate at 30°C and 220 rpm for 2 days. Cells were washed by centrifugation (3,000 g; 5 min.) and resuspension in dH_2_O to OD_600_=0.3-0.5. To assess whether yeast can metabolise RBH and BABA, 5 uL of the suspension was added to 2 mL Yeast Nitrogen Base and increasing concentrations of RBH or BABA (0.2 - 5 mM). To assess toxicity of RBH and BABA, 5 uL of the suspension was added to 2 mL Yeast Nitrogen Base with 10 mM ammonium sulfate and increasing concentrations of RBH or BABA (0.2 - 5 mM). To assess competition between L-Ala and RBH or BABA, 5 uL of the suspension was added to 2 mL Yeast Nitrogen Base supplemented with 1 mM L-Ala and increasing concentrations of RBH or BABA (0.2 - 5 mM). Cells were cultivated at 30°C and 220 rpm for 3 days, after which the OD_595_ was determined in a plate reader (FLUOstar OPTIMA; BMG LABTECH; Germany).

### Assessment of uptake and inhibition kinetics of IRI1/LHT1 in yeast

Transformed 22Δ10α cells were grown in Yeast Nitrogen Base supplemented with 10 mM ammonium sulfate at 30°C and 220 rpm for 2 days. Yeast cells were collected by centrifugation (3000 g; 5 min), washed in dH_2_O, and resuspended in washing buffer (0.6 M sorbitol, 50 mM sodium phosphate, pH 4.5) to OD_600_ = 5. Before the uptake assay, cells were energised by adding 1 M glucose to the growth medium for 10 min. To assess time-dependent uptake of L-[^14^C]Ala in EV- and *IRI1/LHT1*-transformed cells (Supplementary Fig. 6), 1.5-mL of the energised cell culture was added to 1.5 mL uptake buffer, containing 50 nCi L-[^14^C]Ala (158 mCi/mmol; Perkin Elmer; NEC856) with unlabelled L-Ala (50 or 500 μM). After 2, 5 and 10 min. of incubation in a thermomixer (Grant bio ES-20; Grant Instruments; UK; 30°C, 220 rpm), the cell suspensions were mixed with 2 mL ice-cold water and kept on ice to inhibit L-Ala uptake. Cells were then centrifuged (3000 g; 5 min) and washed 4 times with 2 mL ice-cold water, after which pellets were stored at −20 °C for quantification of radioactivity the following day. To quantify uptake and inhibition kinetics (Figs. 6E-F), *IRI1/LHT1*-transformed cells were incubated in the same uptake medium, containing 50 nCi L-[^14^C]Ala with increasing concentrations (1-3000 μM) of unlabelled L-Ala and/or 500 μM inhibitory RBH or BABA. After 5 min of incubation, cells were washed, collected and stored as described above. To assess radioactivity, frozen pellets were resuspended in 750 μL dH_2_O from which 200 μL was loaded onto Combusto-Pad (Perkin Elmer, part number 5067034), which were combusted in a sample oxidiser (Model 307 Sample Oxidizer; Perkin Elmer; USA). Trapped ^14^CO_2_ was quantified by liquid scintillation counting (Tri-Carb 3100TR; Perkin Elmer; USA). L-Ala uptake values over the 5 min. time window were expressed as fmol L-Ala/cell and plotted against the L-Ala concentration to calculate Km and Ki values, using the R package ‘drc’(Ritz et al., 2015).

## ACKNOWLEDGEMENTS

We thank Dr. Henrik Svennerstam for providing the seeds of the *Pro35S:IRI1/LHT1* line, Professor Guillaume Pilot for providing 22Δ10α yeast line, Professor Stephen Rolfe and Dr. Pedro Rocha for advice on the enzyme kinetic experiment, and thank Dr. Karin Posthuma (Enza Zaden) for advice and support throughout the project. This work was supported by a grant from the European Research Council (ERC; no. 309944 “*Prime-A-Plant*”) to J.T., a Research Leadership Award from the Leverhulme Trust (no. RL-2012-042) to J.T., a BBSRC-IPA grant to J.T. (BB/P006698/1) and supplementary grant from Enza Zaden to J.T., and a ERC-PoC grant to JT (no. 824985 *“ChemPrime*). The authors declare no financial conflict of interest.

## AUTHOR CONTRIBUTIONS

J.T. conceived the research; C.-N.T, W.B., P.Z., R.S., and J.T. designed the experiments; C.-N.T, W.B., P.Z., H.W., and I.J. conducted the experiments; C.-N.T, W.B., P.Z., and J.T. analysed the data; C.-N.T, W.B., and J.T. wrote the paper.

## Supplemental Material

**Supplemental Figure 1. Validation of putative *iri* mutants at stage 3 of the mutant screen.**

**(A)** Experiment to confirm statistical robustness of the bioassay system used to verify putative *iri* mutants. The results demonstrate uniformity of *Hpa* colonisation between 20 independent batches of naïve (control-treated) plants and between 20 independent batches of plants expressing RBH-IR (right). Shown are levels of *Hpa* colonisation in leaves of water-treated (left) and RBH-treated (right) plants (Col-0) within one experiment. Each batch consisted of one 60-mL pot containing 20-30 seedlings. Two-week-old seedlings were soil-drenched with water (control) or RBH (0.5 mM) and harvested for trypan-blue staining at at 6 dpi (n = 60-80 leaves) and microscopically assigned to four different classes of *Hpa* colonisaiton (see Fig. **1A** for details). Robustness of the bioassay assay system is indicated by the lack of statistically significant differences in *Hpa* colonisation between independent batches of the same treatment (Fisher’s Exact Test,*p* < 0.05).

**(B)** Quantification of RBH-IR against *Hpa* in 427 putative *iri* lines, using the RBH-IR bioassay. The red y-axis on the right plots *p*-values of the difference in *Hpa* colonisation between RBH-treated mutant plants and RBH-treated Col-0 plants within each sub-experiment (Fisher’s Exact Test), indicating loss of RBH-IR. The dotted line indicates the trehshold of significance (*p* = 0.05).

**Supplemental Figure 2. Characterisation of RBH-IR in mutants carrying independent T-DNA insertions in the SALK/SAIL-annotated genes of *iri2-1, iri 3-1* and *iri4-1*.**

**(A)** Models of T-DNA tagged genes. Red triangles indicate independent T-DNA insertions in the annotated genes of *iri2-1, iri 3-1* and *iri4-1*.

**(B)** Quantification of RBH-IR against *Hpa* in leaves of mutants carrying the T-DNA insertion indicated above. Two-week-old seedilngs were soil-drenched with 0.5 mM RBH and challenge with *Hpa* conidiospores 2 days later. Shown are frequency distributions of trypan-blue-stained leaves across the four *Hpa* colonisation classes (Fig. 1A) at 7 dpi. Asterisks indicate statistically significant differences between control- and RBH-treated samples (Fisher’s exact tests; p < 0.05; n=70-80 leaves).

**(C)** Quantification of arrested *Hpa* colonisation by callose at 3 dpi. *Hpa*-induced callose was analysed in aniline blue/calcofluor-stained leaves by epifluorescence microscopy. Shown are percentages of callose-arrested and non-arrested conidiospores as detailed by Schwarzenbacher et al. (2020). Different letters indicate statistically significant differences in frequencies between control- and RBH-treated plants (Fisher’s exact tests; *p* < 0.05; n>100 conidiospores).

**Supplemental Figure 3. Genetic characterisation of two independent *iri1/lht1* mutant lines and the *IRI1/LHT1* over-expression line.**

**(A)** Gene model of *IRI1/LHT1*. Red triangles indicate T-DNA insertions in the *iri1-1/lht1-5* and *iri1-2/lht1-4* mutant. Dark blue arrows indicate aligned locations of the primers used for RT-qPCR analysis (see below).

**(B)** Confirmation of T-DNA insertions by PCR. Shown are PCR products of the expected sizes in Col-0, *iri1-1/lht1-5* and *iri1-2 /lht1-4*, using gene-specific primers (LP+RP) and the left-border primer of the T-DNA in combination with the gene-specific RP primer (LBb1.3+RP). Primer sequences are listed in Supplemental Table 1.

**(C)** RT-qPCR quantification of IRI1/LHT1 gene expression in Col-0 and the over-expression line *Pro35S:IRI1/LHT1*. Biologically replicated seedling samples (n=5) were taken from 2-week-old seedlings under the same growth conditions as our IR assay. Boxplots show median (middle bar), interquartile range (IQR; box), 1.5 x IQR (whiskers) and replication units (single dots) of relative expression values normalised to the mean relative expression value of Col-0.

**Supplemental Figure 4. Transgenic over-expression of *IRI1/LHT1* improves Arabidopsis growth on medium with L-Alanine as only N source, which is antagonised by co-application of RBH.**

**(A)** Shown are 2-week-old seedlings of Col-0 (upper left), *iri1-1/iht1-5* (upper right), and *Pro35S:IRI1/LHT1* (bottom) on MS agar plates without inorganic N and supplemented with increasing concentrations of RBH and/or L-Alanine. All genotypes failed to grow on medium with RBH as only N source.

**(B)** Green leaf areas (GLA± SEM; n=10-19) of 2-week-old Col-0, *iri1-1/iht1-5*, and *Pro35S:IRI1/LHT1* seedlings from the same experiment. Inset shows p-values of effects on green leaf area by RBH concentration, L-Alanine concentration and interaction thereof inside each genotype (2-way ANOVA).

**Supplemental Figure 5. RBH and BABA have minimal effects on yeast growth but cannot be used as N source by yeast**.

The yeast mutant strain 22Δ10α (Besnard et al., 2016) was transformed with the Arabidopsis *IRI1/LHT1* gene and grown in liquid growth medium with (top) or without (NH_4_)_2_SO_4_ (bottom), co-supplied with increasing concentration of RBH (left) or BABA (right). Planktonic growth was quantified spectrophotometrically by determining ΔOD_595_ values (OD_595_ - OD_595_ of medium without yeast). Shown by average ΔOD_595_ values (±SME; n=4) after growth for 3 days.

**Supplemental Figure 6. Transformation of the 22Δ10α mutant of yeast with the *IRI1/LHT1* gene complements uptake of L-[^14^C] Alanine.** Empty vector (EV)- and *IRI1/LHT1-transformed* cells were incubated for 2, 5 and 10 min. in the presence of 50 or 500 μM L-Ala with 50 nCi ^14^C-labelled L-Ala. Data represent average cellular L-Ala concentrations (fmol L-Ala/cell; ±SEM; n=3).

**Supplemental Data Set 1.** Annotated genomic T-DNA insertions of the 108 confirmed *iri* lines, RBH-IR phenotypes, and expression profiles of the associated T-DNA-tagged genes.

**Supplemental Table 1.**
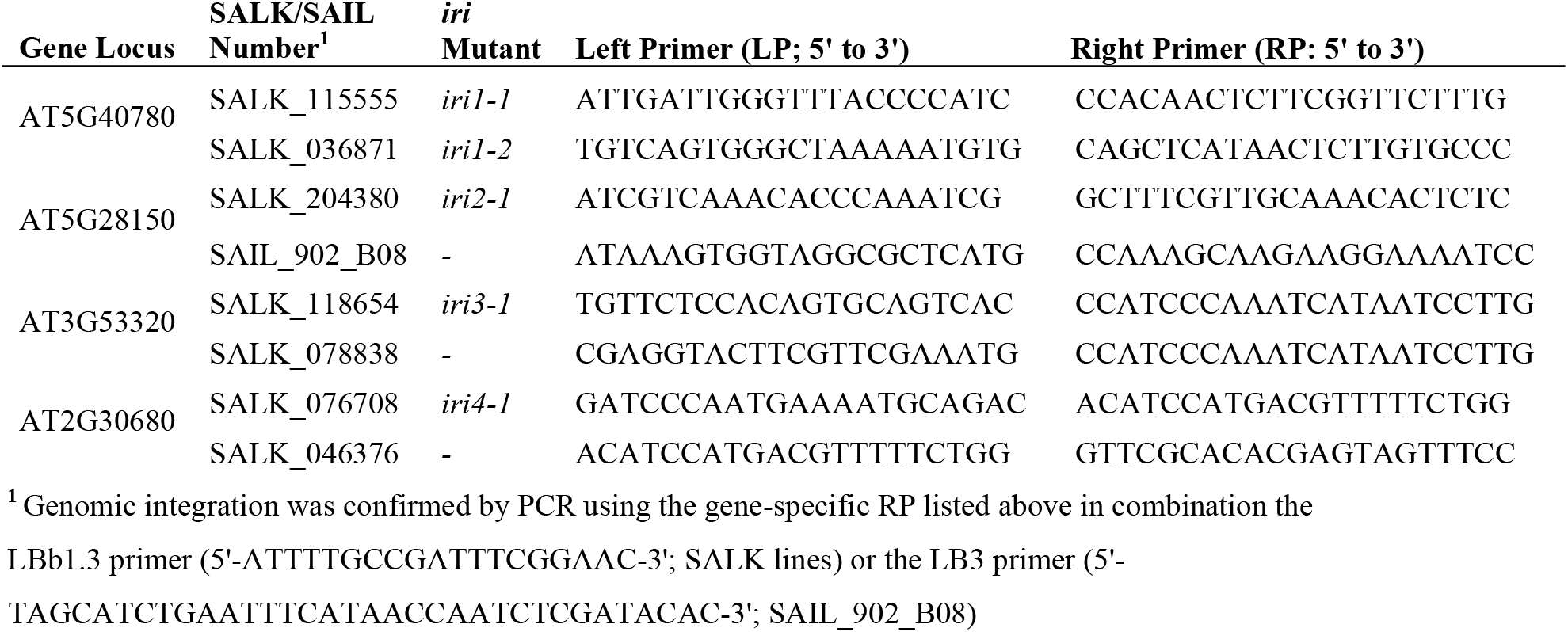
Genomic primers used for characterisation of T-DNA insertion lines

